# The Functional False Discovery Rate with Applications to Genomics

**DOI:** 10.1101/241133

**Authors:** Xiongzhi Chen, David G. Robinson, John D. Storey

**Affiliations:** Lewis-Sigler Institute for Integrative Genomics, Princeton University, Princeton, NJ 08544, USA; Present address: Department of Mathematics and Statistics, Washington State University, Pullman, WA 99163, USA

## Abstract

The false discovery rate measures the proportion of false discoveries among a set of hypothesis tests called significant. This quantity is typically estimated based on p-values or test statistics. In some scenarios, there is additional information available that may be used to more accurately estimate the false discovery rate. We develop a new framework for formulating and estimating false discovery rates and q-values when an additional piece of information, which we call an “informative variable”, is available. For a given test, the informative variable provides information about the prior probability a null hypothesis is true or the power of that particular test. The false discovery rate is then treated as a function of this informative variable. We consider two applications in genomics. Our first is a genetics of gene expression (eQTL) experiment in yeast where every genetic marker and gene expression trait pair are tested for associations. The informative variable in this case is the distance between each genetic marker and gene. Our second application is to detect differentially expressed genes in an RNA-seq study carried out in mice. The informative variable in this study is the per-gene read depth. The framework we develop is quite general, and it should be useful in a broad range of scientific applications.

## 1 Introduction

Multiple testing is now routinely conducted in many scientific areas. For example, in genomics, RNA-seq technology is often utilized to test thousands of genes for differential expression among two or more biological conditions. In expression quantitative trait loci (eQTL) studies, all pairs of genetic markers and gene expression traits can be tested for associations, which often involves millions or more hypothesis tests. The false discovery rate (FDR, Benjamini and Hochberg, 1995) and the q-value (Storey, 2002, 2003) are often employed to determine significance thresholds and quantify the overall error rate when testing a large number of hypotheses simultaneously. Therefore, improving the accuracy in estimating false discovery rates and q-values remains an important problem.

In many emerging applications, additional information on the status of a null hypothesis or the power of a test may be available to help better estimate the FDR and q-value. For example, in eQTL studies, gene-SNP^1^ basepair distance informs the prior probability of association between a gene-SNP pair, with local associations generally more likely than distal associations (Brem et al., 2002; Ronald et al., 2005; Doss et al., 2005). A second example comes from RNA-seq studies, for which per-gene read depth informs the statistical power to detect differential gene expression (Tarazona et al., 2011; Cai et al., 2012) or the prior probability of differential gene expression (Robinson et al., 2015). Genes with more sequencing reads mapped to them (i.e., higher per-gene read depth) have greater ability to detect differential expression or may be more likely to be differentially expressed than do low depth genes.

Figure 1 shows results from multiple testing on a genetics of gene expression study (Smith and Kruglyak, 2008) and an RNA-seq differential expression study (Bottomly et al., 2011). In the genetics of gene expression study, the p-values are subdivided according to six different gene-SNP basepair distance strata. In the RNA-seq study, the p-values are subdivided into six different strata of per-gene read depth. It can be seen in both cases that the proportion of true null hypotheses and the power to identify significant tests vary in a systematic manner across the strata. The goal of this paper is to take advantage of this phenomenon so that we may improve the accuracy of calling tests significant and do so without having to create artificial strata as in Figure 1.

**Figure 1:**
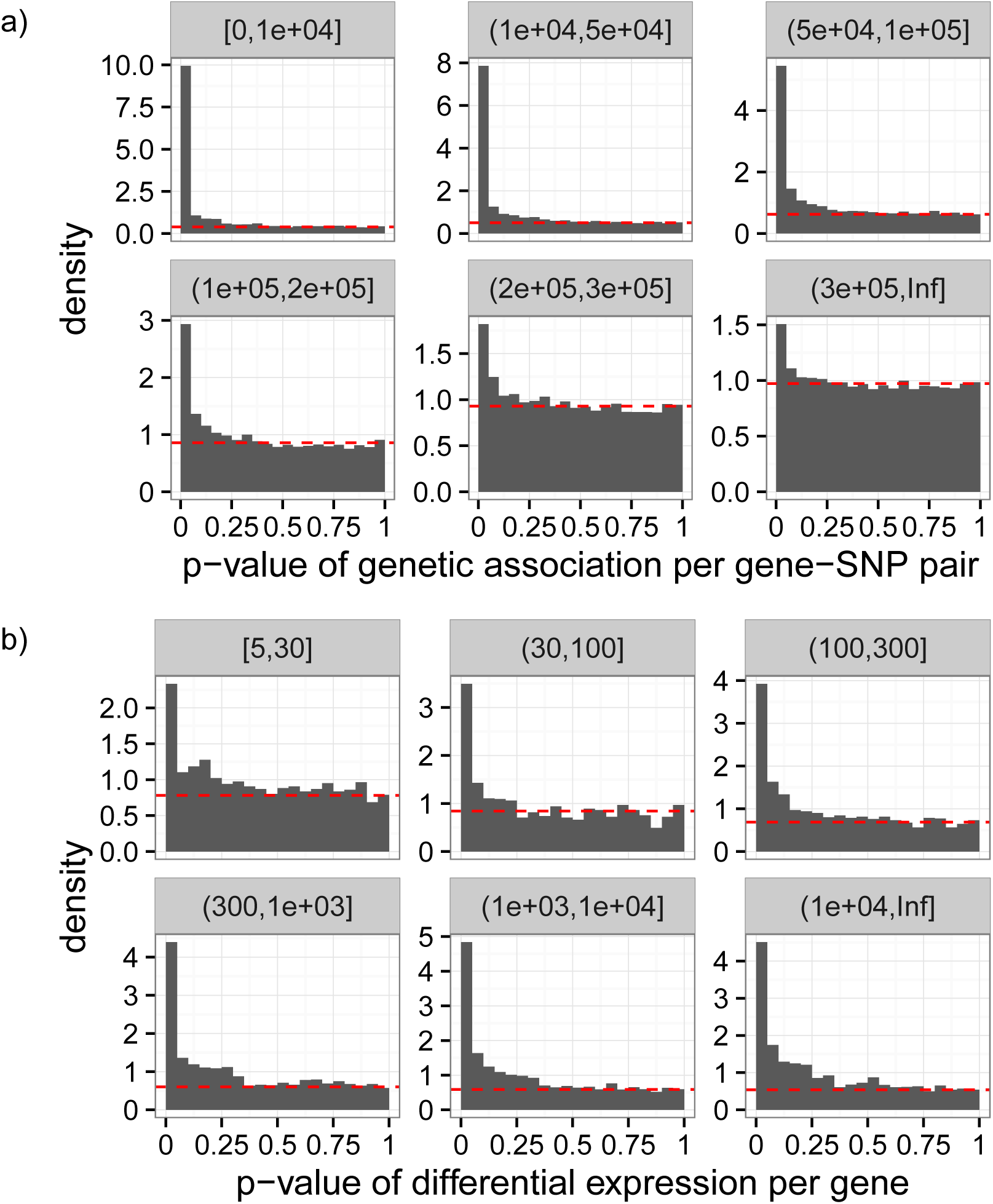
a) P-value histograms of Wilcoxon tests for genetic association between genes and SNPs for the eQTL experiment in Smith and Kruglyak (2008), divided into six strata based on the gene-SNP basepair distance indicated by the strip names. The null hypothesis is “no association between a gene-SNP pair”. b) P-value histograms for assessing differential gene expression in the RNA-seq study in Bottomly et al. (2011), divided into six strata based on per-gene read depth indicated by the strip names. The null hypothesis is “no differential expression (for a gene) between two conditions”. In each subplot, the estimated proportion of true null hypotheses for all hypotheses in the corresponding stratum is based on Storey (2002) and indicated by the horizontal dashed line. It can be seen that gene-SNP genetic distance or per-gene read depth provides affects the prior probability of a gene-SNP association or differential gene expression.

We propose the *functional FDR* (fFDR) methodology that efficiently incorporates additional quantitative information for estimating the FDR and q-values. Specifically, we code additional information into a quantitative *informative variable* and extend the Bayesian framework for FDR pioneered in Storey (2003) to incorporate this informative variable. This leads to a functional proportion of true null hypotheses (or “functional null proportion” for short) and a functional power function. From this, we derive the optimal decision rule utilizing the informative variable. We then provide estimates of functional FDRs, functional local FDRs, and functional q-values to utilize in practice.

Related ideas have been developed, such as p-value weighting (Genovese et al., 2006; Roquain and van de Wiel, 2009; Ignatiadis et al., 2016), stratified FDR control (Sun et al., 2006), stratified local FDR thresholding (Ochoa et al., 2015), and covariate-adjusted proportion of true null hypotheses estimation Boca and Leek (2017). Stratified FDR and local FDR rely on clearly defined strata, which may not always be available or make the best use of information. Covariate-adjusted proportion of true null hypotheses estimation focuses on only one component of the FDR. P-value weighting has been a successful strategy, but it remains challenging to derive weights that indeed result in improved power subject to a target FDR level. In particular, how to obtain optimal weights under different optimality criteria is still an open problem.

Our methodology serves as an alternative to p-value weighting. We are motivated by similar scientific applications as the “independent hypothesis weighting” (IHW) method proposed in Ignatiadis et al. (2016), which employs a covariate to improve the power of multiple testing. Our approach is distinct from this since IHW is a weighted version of the Benjamini-Hochberg procedure (Benjamini and Hochberg, 1995). We work with a Bayesian model developed in Storey (2003), providing direct calculations of optimal significance thresholds and take an empirical Bayes strategy to estimate several FDR quantities. As earlier frequentist and Bayesian approaches to the FDR in the standard scenario both proved to be important, our contribution here should serve to complement the p-value weighting strategy.

To demonstrate the effectiveness of our proposed methodology, we conduct simulations and analyze two genomics studies, an RNA-seq differential expression study and a genetics of gene expression study. In doing so, we uncover important operating characteristics of the fFDR methodology and we also provide several strategies for visualizing and interpreting results. Although our applications are focused on genomics, we anticipate the framework presented here will be useful in a wide range of scientific problems.

This article is organized as follows. We formulate the fFDR methodology in Section 2 and provide its implementation in Section 3. A simulation study on the fFDR methodology is provided in Section 4, and two applications of the methodology are given in Section 5. We end the article with a discussion in Section 6.

## 2 The functional FDR framework

In this section, we formulate the fFDR theory and methodology. To this end, we first introduce our model, provide formulas for the positive false discovery rate (pFDR), positive false nondiscovery rate (pFNR) and q-value, and then describe the significance rule based on the q-value.

### 2.1 Joint model for p-value, hypothesis status, and informative variable

Let *Z* be the informative variable and assume it has been transformed to be uniformly distributed on the interval [0, 1] so that *Z* ~ Uniform (0, 1). For example, *Z* can denote the quantiles of the per-gene read depths in an RNA-seq study or the quantiles of the genomic distances in an eQTL experiment. Denote the status of the null hypothesis by *H*, such that *H* = 0 when the null hypothesis is true and *H* = 1 when the alternative hypothesis is true. We assume that conditional on *Z* = *z* the null hypothesis is *a priori* true with probability *π*_0_ (*z*), i.e.,

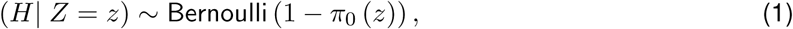
where the function *π*_0_(*z*) ranges in [0, 1]. We call *π*_0_(*z*) the “prior probability of the null hypothesis”, “functional proportion of true null hypotheses”, or “functional null proportion.” When *π*_0_(*z*) is constant, it will be simply denoted by *π*_0_.

To formulate the distribution of the p-value, *P*, we assume the following: (i) when the null hypothesis is true, (*P*|*H* = 0, *Z*) ~ Uniform(0,1) regardless of the value of *Z*; (ii) when the null hypothesis is false, the conditional density of (*P*|*H* = 1, *Z* = *z*) is *f*_1_(·|*z*). The conditional density of *P* given *Z* = *z* is then

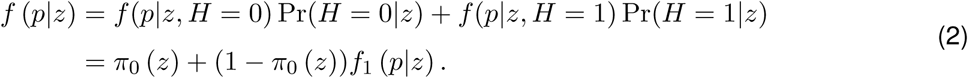

Since *Z* has constant density *f*(*z*) = 1, the joint density *f*(*p*, *z*) = *f*(*p*|*z*) for all (*p*, *z*) ∈ [0, 1]^2^ so that

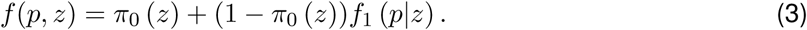

The transformation of *Z* into the Uniform(0,1) distribution and the representation in equation (3) is important because it is more straightforward to estimate *f*(*p*, *z*) than *f*(*p*|*z*)

### 2.2 Optimal statistic

Now suppose there are *m* hypothesis tests, with *H_i_* for *i* = 1, 2,…, *m* indicating the status of each hypothesis test as above. For example, *H_i_* can denote whether gene *i* is not differentially expressed or not, or whether there is an association between the *i*th gene-SNP pair. For the *i*th hypothesis test, let its calculated p-value be *p_i_* and its measured informative variable be *z_i_*; additionally let *P_i_* and *Z_i_* be their respective random variable representations.

Let *T* = (*P*, *Z*) and *T_i_* = (*P_i_,Z_i_*) for 1 ≤ *i* ≤ *m*. Suppose the triples (*T_i_*, *H_i_*) are independent and identically distributed (i.i.d.) as (*T*, *H*). If the same significance region Γ in [0, 1]^2^ is used for the *m* hypothesis tests, then identical arguments in the proof of Theorem 1 in Storey (2003) imply that pFDR(Γ) = Pr(*H* = 0|*T* ∈ Γ) and pFNR(Γ) = Pr(*H* = 1|*T* ∉Γ), where pFDR is the positive false discovery rate and pFNR is the false non-discovery rate as defined in Storey (2003). The bivariate function

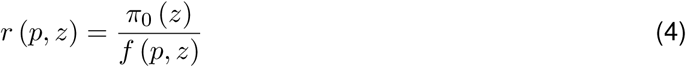
defined on [0, 1]^2^ is the posterior probability that the null hypothesis is true given the observed pair (*p*, *z*) of p-value and informative variable. Note that *r*(*p*, *z*) = Pr(*H* = 0|*T* =(*p*, *z*)), so this quantity is an extension of the local FDR (Storey, 2003; Efron et al., 2001) also known as the posterior error probability (Kall et al., 2008). Straightforward calculation shows that

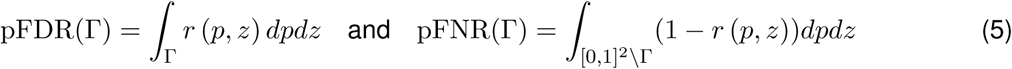

Define significance regions {Γ*_τ_*: *τ* ∈ [0, 1]} with

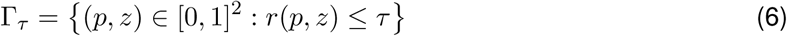
such that test *i* is statistically significant if and only if *T_i_* ∈ Γ*_τ_*. Then by identical arguments in Section 6 leading up to Corollary 4 in Storey (2003), Γ*_τ_* gives the Bayes rule for the Bayes error

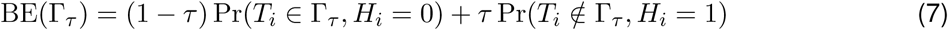
for each *τ* ∈ [0, 1]. Therefore, by arguments analogous to those in Storey (2003), *r*(*p*, *z*) is the optimal statistic for the Bayes rule with Bayes error (7).

### 2.3 Q-value based decision rule

With the statistic *r*(*p*, *z*) in (4) and nested significance regions {Γ*_τ_*: *τ* ∈ [0, 1]} with Γ*_τ_* defined by (6), the definition of q-value in Storey (2003) implies that the q-value for the observed statistic *t* = (*p*, *z*) is

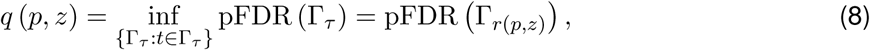
where the second equality follows from Theorem 2 in Storey (2003), noting that {Γ*_τ_*} are constructed from the posterior probabilities *r*(*p*, *z*). Estimating the q-value in (8) will be discussed in Section 3.3.

Let *q*(*p_i_*, *z_i_*) denote the q-value of *t_i_* = (*p_i_*, *z_i_*) for the *i*th null hypothesis *H_i_*. At a target pFDR level *α* ∈ [0, 1], the following signficance rule

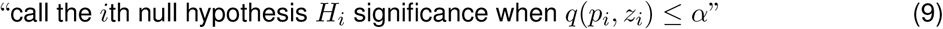
has pFDR no larger than *α*. Note that this significance rule is identical to the significance regions {Γ*_τ_*}from above, so significance rule (9) also achieves the Bayes optimality for the loss function (7). We refer to (9) as the “Oracle”, which will be estimated by a procedure detailed below in Section 3. When only p-values 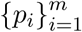 are used, the above significance rule becomes

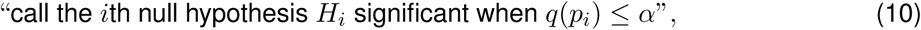
where *q*(*p_i_*) is the original q-value for *p_i_* as developed in Storey (2002) and Storey (2003).

## 3 Implementation of the fFDR methodology

We aim to implement the decision rule in (9) for the fFDR methodology by a plug-in estimation procedure. For this, we need to estimate the two components of the statistic *r*(*p*, *z*) given in (4): the functional null proportion *π*_0_(*z*) and the joint density *f*(*p*, *z*) with support on [0, 1]^2^. We also need to estimate the q-value defined in (8). We will provide in Section 3.1 three complementary methods to estimate *π*_0_(*z*), in Section 3.2 a kernel-based method to estimate *f*(*p*, *z*), and in Section 3.3 the estimation of q-values and the plug-in procedure.

### 3.1 Estimating the functional null proportion

Our proposed approaches to estimate the functional null proportion *π*_0_(*z*) is based on an extension of the approach taken in Storey (2002). Recalling that *π*_0_(*z*) = Pr(*H* = 0|*Z* = *z*), it follows that for each *λ* ∈ [0, 1),

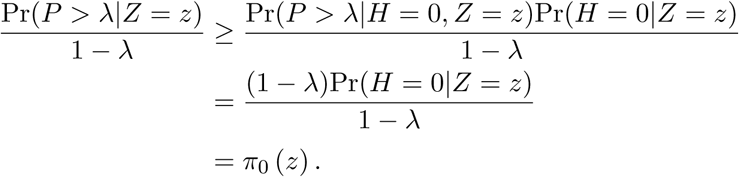

If we define the indicator function *ξ_λ_*(*z*)=1_{_*_P>λ_*_|_*_Z_*_=_*_z_*_}_, then

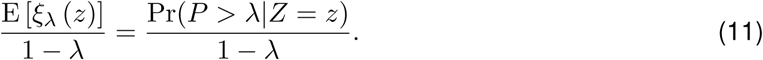

Therefore, E[*ξ_λ_* (*z*)]/(1−*λ*) is a conservative estimate of *π*_0_(*z*) and it will form the basis of our estimate of *π*_0_(*z*).

Our first method to estimate *π*_0_(*z*) is referred to as the “GLM method” since it estimates E[*ξ_λ_* (*z*)] using generalized linear models (GLM). For each *z* ∈ [0, 1], we let *η*(*z*) = *β*_0_ + *β*_1_*z* and

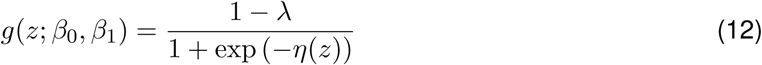
for two parameters (*β*_0_*,β*_1_), and then fit

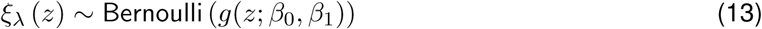
to obtain an estimate 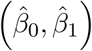 of (*β*_0_*,β*_1_) using the paired realizations 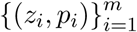. We then estimate *π*_0_(*z*) by

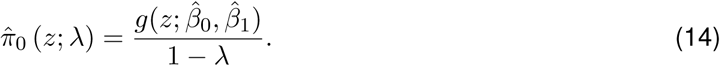

Our second method to estimate *π*_0_(*z*) is referred to as the “GAM method” since it estimates E[*ξ_λ_* (*z*)] using generalized additive models (GAM). Specifically, we model *η* in the GLM method as a nonlinear function of *z* via GAM while keeping the functional form of *g* in (12) and the estimator (14). For example, η can be modeled by B-splines (Hastie and Tibshirani, 1986; Wahba, 1990) whose degree can be chosen, e.g., by generalized cross-validation (GCV) (Craven and Wahba, 1978). The GAM method removes the restriction induced by the GLM method that *π*_0_(*z*) be a monotone function of *z*.

Our third method to estimate *π*_0_(*z*) is referred to as the “Kernel method” since it estimates E[*ξ_λ_* (*z*)] via kernel density estimation (KDE). Since *Z* ~ Uniform (0, 1), it follows that

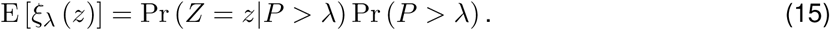

To estimate E[*ξ_λ_* (*z*)], we estimate the two factors in the right hand side of (15) separately. It is straight-forward to see that the estimator from Storey (2002),

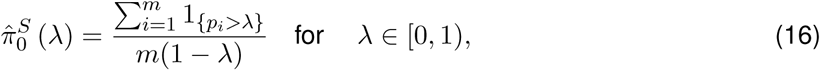
is a conservative estimator of Pr(*P* > *λ*)/(1 − *λ*). Further, if we let 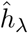 be a conservative estimator of the density of the *z_i_*’s whose corresponding p-values are greater than *λ*, then 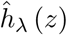 conservatively estimates Pr (*Z* = *z*|*P* > *λ*). Correspondingly,

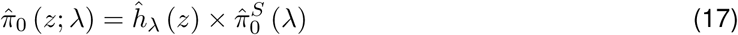
is a conservative estimator of *π*_0_(*z*). In the implementation, we obtain 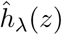 using the methods in Geenens (2014) since *z* ranges in the unit interval. Note that (17) is essentially a nonparametric alternative to (14) since 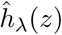 does not have the constraint on its shape that 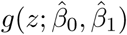 does.

To maintain a concise notation, we write 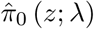 as 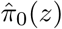. If no information on the shape of *π*_0_(*z*) is available, we recommend using the Kernel or GAM method to estimate *π*_0_(*z*); if *π*_0_(*z*) is monotonic in *z*, then the GLM method is preferred. An approach for automatically handling the tuning parameter *λ* for the estimators is provided in Appendix A.

### 3.2 Estimating the joint density *f*(*p*, *z*)

The estimation of the joint density *f*(*p*, *z*) of the p-value *P* and informative variable *Z* involves two challenges: (i) *f* is a density function defined on the compact set [0, 1]^2^; (ii) *f*(*p*, *z*) may be monotonic in *p* for each fixed *z*, requiring its estimate to also be monotonic. In fact, in the simulation design below in Section 4.1, *f*(*p*, *z*) is monotonic in *p*. To deal with these challenges, we estimate *f* in a two-step procedure as follows. Firstly, to address the challenge of density estimation on a compact set, we use a local likelihood KDE method with a probit density transformation (Geenens, 2014) to obtain an estimate 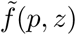 of *f*(*p*, *z*), where an adaptive nearest-neighbor bandwidth is chosen via GCV. Secondly, if *f*(*p*, *z*) is known to be monotonic in *p* for each fixed *z*, then we utilize the algorithm in Appendix B to produce an estimated density 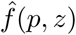 that has the same monotonicity property as *f*(*p*, *z*) at the observations 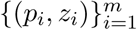.

### 3.3 FDR and q-value estimation

With the estimates 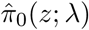 and 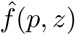 respectively for *π*_0_(*z*) and *f*(*p*, *z*), the functional posterior error probability (or local FDR) statistic *r*(*p*, *z*) in (4) is estimated by

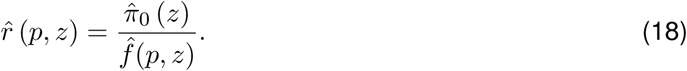

For a threshold *τ*, Storey et al. (2005) proposed the following pFDR estimate

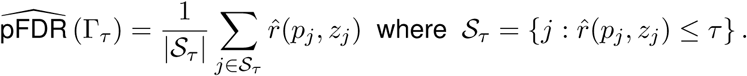

The rationale for this estimate is that the numerator is the expected number of false positives given the posterior distribution 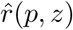 and the denominator is the expected number of total discoveries given 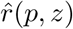 (which is directly observed). This is related to a semiparametric Bayesian procedure detailed in Newton et al. (2004). Given this, the functional q-value *q*(*p_i_*, *z_i_*) of *t_i_* =(*p_i_*, *z_i_*) corresponding to the *i*th null hypothesis *H_i_* is estimated by:

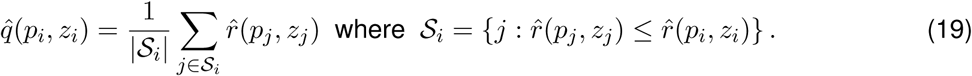

The plug-in decision rule is to call the null hypothesis *H_i_* significant whenever 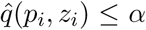 at a target pFDR level *α*. In this work, we refer to this rule as the “functional FDR (fFDR) method”.

Recall that an estimate of 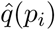 of the q-value *q*(*p_i_*) for *p_i_* can be obtained by the q-value package (Storey et al., 2015). Then the plug-in decision rule based on 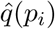 is to call the null hypothesis *H_i_* significant whenever 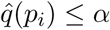. This rule is referred to as the “standard FDR method” in this work.

## 4 Simulation study

We now present a simulation study on the performance of the fFDR method. To assess the overall error rate of a multiple testing procedure (MTP), we compare the estimated q-values to the oracle q-values. To assess the power of an MTP, we use the expected ratio of the number of significant true alternative hypothesis tests to the total number of true alternative tests, which is the average power across all true alternative tests. Note that the “oracle q-value” and “average power” are calculated as Monte Carlo averages over the simulated data sets for each scenario based on the fact that we know the true status of each hypothesis test.

### 4.1 Simulation design

To investigate the performance of the fFDR method when the functional null proportion *π*_0_(*z*) takes different forms, we construct four types of *π*_0_(*z*), where we calculate the average value as 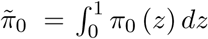:

1. *decreasing*: *π*_0_(*z*)=1 − (0.9 × *z*^3.5^ + 0.01) such that 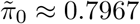.
2. *increasing*: *π*_0_(*z*)=0.9 × (*z*^0.3^ + 0.1) such that 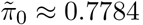.
3. *constant*: *π*_0_ (*z*) = *π*_0_ = 0.85 for all values of *z*.
4. *sine*: *π*_0_(*z*) = 0.2 + 0.78 × sin(*πz*) such that 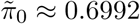.

To investigate how the performance of the fFDR method is affected by the informative variable *Z* through the conditional p-value density *f*_1_(*p*|*z*) under the false null hypothesis, we construct two types of *f*_1_(*p*|*z*):

1. *dependent*: 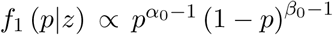, where *α*_0_ = 0.3, *γ*_0_ = 4.5, *β*_0_ = *γ*_0_ + 1.4*z*, where the scalar 1.4 helps reduce the chance of generating large p-values under the false null hypothesis.
2. *independent*: 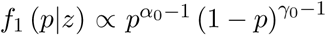 with *α*_0_ = 0.3, *γ*_0_ = 4.5, where *f*_1_ (*p*|*z*) is independent from *z*.

The choices of *π*_0_(*z*) are based on our observations from real data sets that *π*_0_ (*z*) infrequently assumes the value 0 or 1, and that the mean of *π*_0_ (*Z*) is not very small. Note that *f*_1_(*p*|*z*) for each fixed *z* is from the Beta density family. With the chosen *α*_0_ and *β*_0_, for each fixed *z* the p-value under the false null hypothesis has a decreasing density, is stochastically smaller than the Uniform(0,1) random variable, and has positive probability to assume a range of small values. This enables the competing MTPs (described in the next paragraph) to have reasonable power. Further, the densities *f*_1_(·|*z*) indexed by *z* for the p-values under the false null hypothesis tend to satisfy *f*_1_(*p*|*z*) ≤ *f*_1_ (*p*|*z*′) for large *p* when *z* ≤ *z*′, and therefore *z* indexes the power of the test statistic that produces the corresponding p-value. The characteristics of *π*_0_(*z*) and *f*_1_(*p*|*z*) are intended to make the simulation study both practical and fair to the competing MTPs.

For each of the eight configurations given above (four versions of *π*_0_(*z*) by two versions of *f*_1_(*p*|*z*)), we simulated 100 data sets for *m* = 3000 tests according to the following procedure.

1. Generate 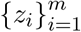 independently from Uniform (0, 1).
2. For each *i* = 1, 2,…, *m*, generate *H_i_*|*z_i_* ~ Bernoulli (1 − *π*_0_ (*z_i_*)); if *H_i_* = 0, generate *p_i_* from Uniform (0, 1) and if *H_i_* = 1 generate *p_i_* according to density *f*_1_(*p*|*z_i_*).
3. Apply the standard FDR method, the fFDR method and the Oracle with q-value cut-offs at 0.01, 0.02,…, 0.1. Calculate the true FDR and average power for each MTP.

### 4.2 Simulation results

We first examined the power and the estimation accuracy for the three methods, where the fFDR method was applied using the GAM model to estimate *π*_0_(*z*). Figure 2 displays the average power with respect to the applied q-value cut-offs. The following can be seen: (i) the power of the fFDR method is very close to that of the Oracle; (ii) the power of the fFDR method is greater than that of the standard FDR method when the functional null proportion *π*_0_(*z*) is nonconstant; (iii) the power of the fFDR method is no smaller than that of the standard FDR method when *π*_0_ is constant. It should be noted that, when *π*_0_(*z*) is nonconstant, the fFDR method achieves ~10% to 30% increase in power over the standard FDR method.

**Figure 2:**
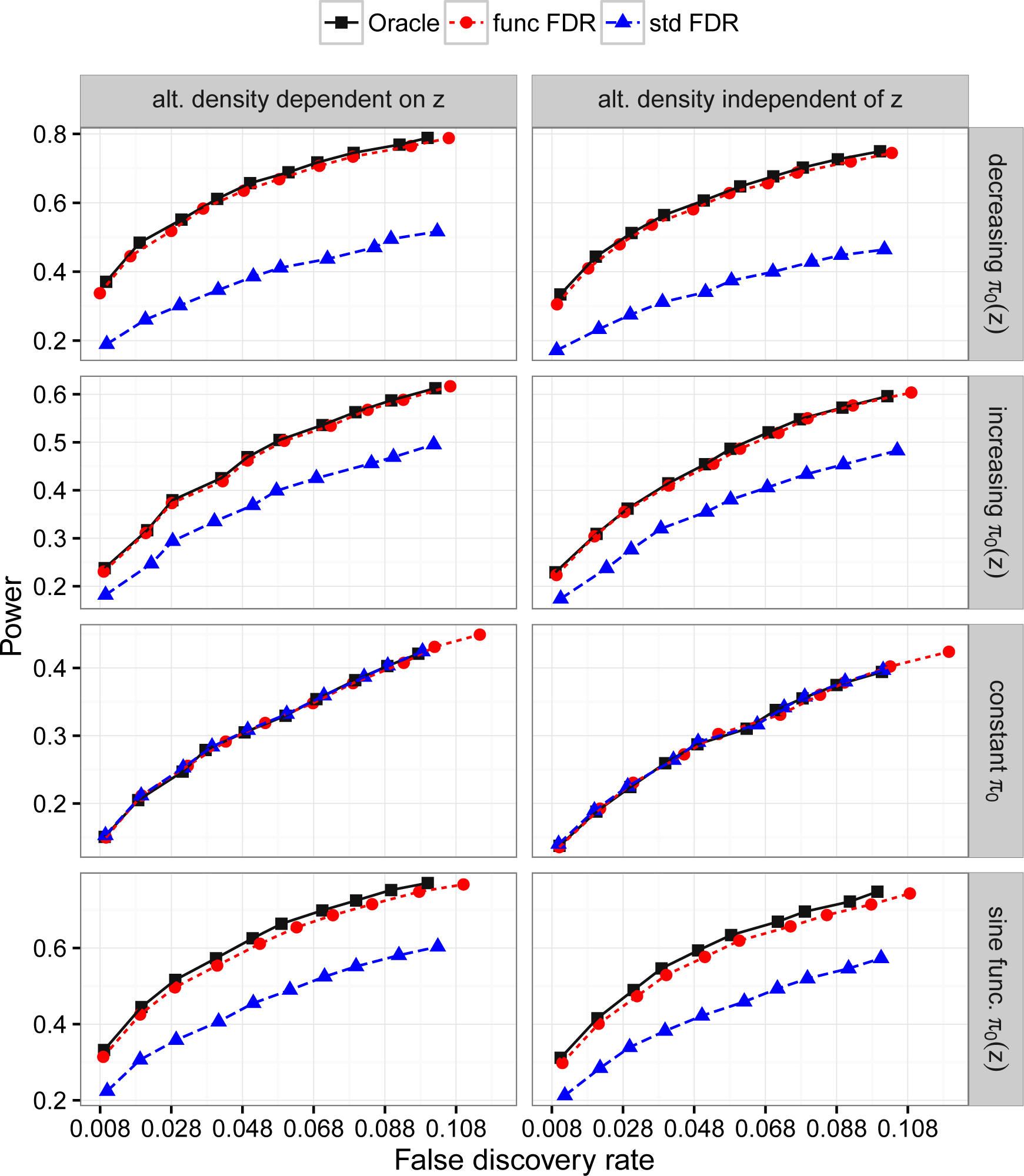
Performance of the fFDR method (func FDR) against that of the standard FDR method (std FDR), with reference to that of the “Oracle”. The points from left to right on each line in each subfigure are successively obtained for q-value cutoffs *α* = 0.01, 0.02,…, 0.1. The column and row plot titles denote the type of *π*_0_(*z*) and alternative density *f*_1_(*p*|*z*) utilized. It can be seen that the power of the fFDR method is very close to that of the Oracle and that it is greater than that of the standard FDR method when *π*_0_(*z*) is not a constant.

Figure 3 shows the estimated q-values versus the oracle q-values, where the oracle q-values are computed using the optimal statistic given in (4). It shows that the fFDR method estimates the oracle q-values accurately and that it does so more accurately than the standard FDR method. In summary, the fFDR method in these simulations is more powerful than the standard FDR method when the functional null proportion is nonconstant and it estimates the FDR accurately. Figure 4 shows the performance of the estimator 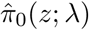 implemented by the GAM, GLM, and Kernel methods. We see that these estimates are accurate but may be slightly downwardly biased in subsets of the range of *π*_0_(*z*). One exception is the GLM estimate in the *sine* scenario, but this is expected given that the GLM model has only a linear term in *z* and is not flexible enough to capture the sine shape. The GLM model could of course be modified to include higher order polynomial terms in *z* to make it more flexible.

**Figure 3:**
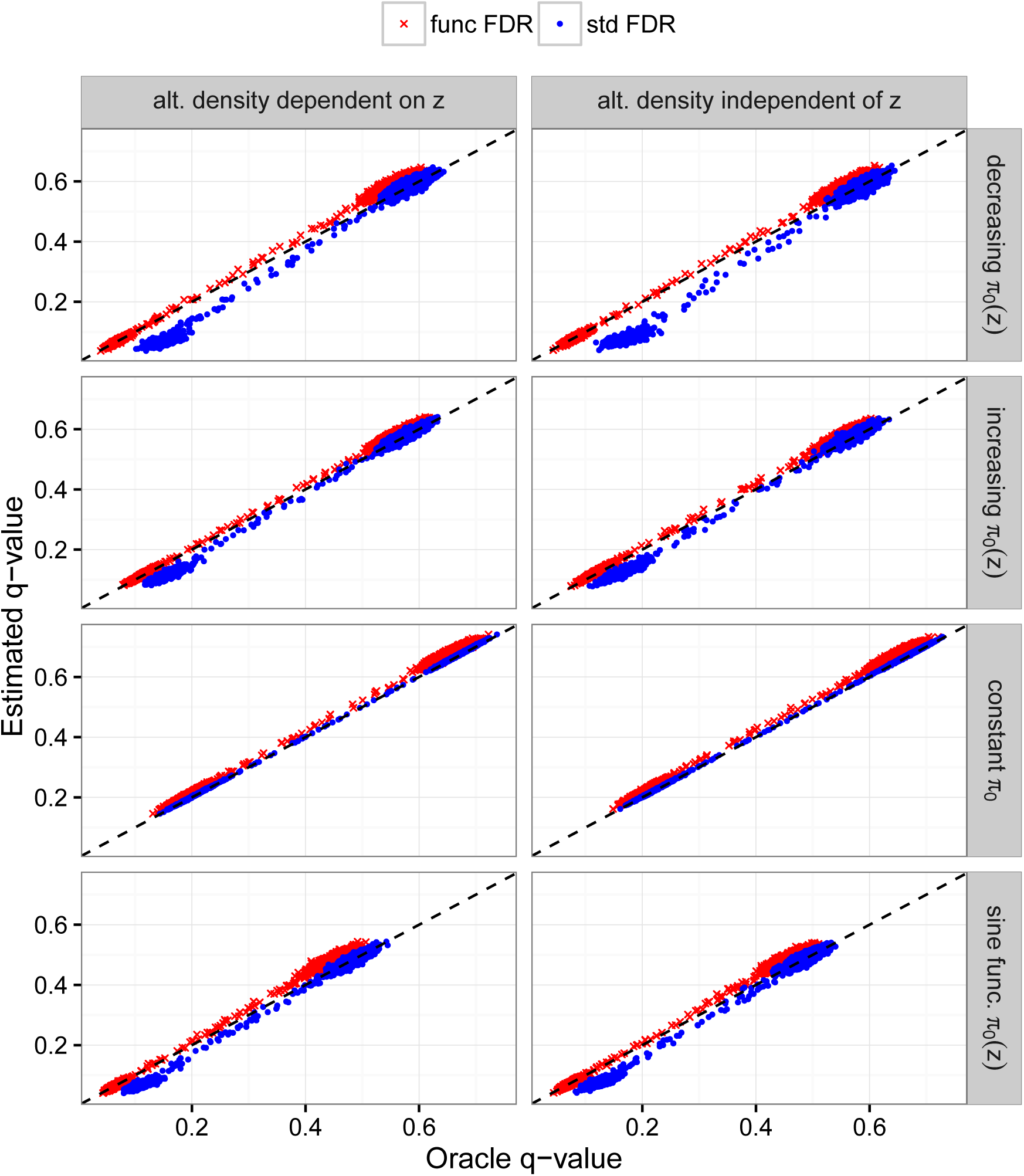
Estimated q-values versus oracle q-values, where the oracle q-values are computed based on the true underlying model. The dashed line in each plot indicates perfect estimates of the oracle q-values. The column and row plot titles denote the type of *π*_0_(*z*) and alternative density *f*_1_(*p*|*z*) utilized. It can be seen that the fFDR method (func FDR) estimates oracle q-values accurately and does so more accurately than the standard FDR method (std FDR).

Figure 2 shows that when *π*_0_ is a constant, there is not an increase in power of the fFDR method over the standard FDR method. We investigated why this might be the case. We first examined the accuracy of estimating *π*_0_ when it is a constant, shown in Figure 5. The GAM and Kernel methods yield similar estimates, whereas the GLM method is almost a constant and very close to the estimate given by the `qvalue` package (Storey et al., 2015). Whereas the GLM, GAM, and Kernel methods yield similar estimates in the *decreasing* and *increasing* scenarios of *π*_0_(*z*) (Figure 4) as well as in the real data analysis shown in the next section (Figure 6), one can see that the GLM method may be more accurate in the case that *π*_0_ is constant. We therefore repeated the simulations for the *constant* scenario where the true *π*_0_ was substituted for the estimated *π*_0_(*z*) in the fFDR method and for the estimated *π*_0_ in the standard FDR method. The power comparisons between fFDR and the standard FDR were qualitatively similar to those observed in the third row of Figure 2.

**Figure 4:**
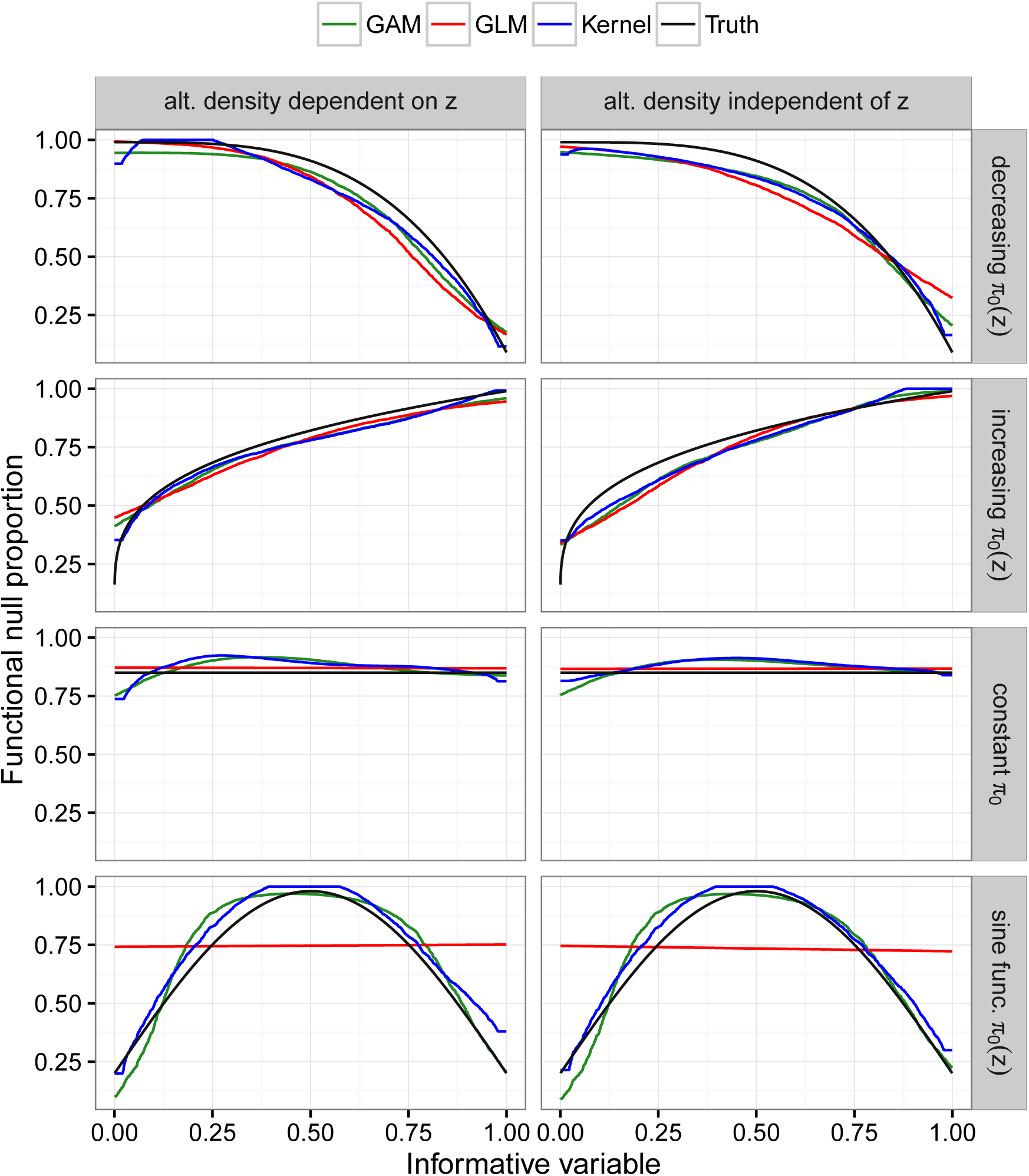
Performance of the estimator 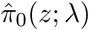 of the functional null proportion *π*_0_(*z*). The column and row plot titles denote the type of *π*_0_(*z*) and alternative density *f*_1_(*p*|*z*) utilized.

This then focused our attention on the estimation of the *f*(*p*, *z*) component from our statistic *r*(*p*, *z*) = *π*_0_(*z*)/*f*(*p*, *z*) of equation (4). Besides the Beta family used in defining *f*(*p*, *z*) in the simulations, we defined *f*(*p*, *z*) from several other distributions (including those induced by Normal and t distributions), but we continued to see similar levels of power between the fFDR method and the standard FDR. An explanation for this is that, when the signal strengths are moderate or strong, the p-values already capture much of the power characteristics of the test statistics so that *f*(*p*, *z*) ≈ *f*(*p*). In this case, the informative variable may not be influential in the *f*(*p*, *z*) component of the *r*(*p*, *z*) statistic. These findings have several relevant implications: (i) if the researcher is certain that *π*_0_(*z*) is constant, then the fFDR method can be implemented with the estimate of a constant *π*_0_; (ii) if the signal strengths are not weak and the fFDR method performs very similarly to the standard FDR method, then it is likely that *π*_0_(*z*) does not depend on the informative variable; (iii) it will be useful to better understand *f*(*p*, *z*) versus *f*(*p*); and (iv) it will be useful to derive a test of whether *π*_0_(*z*) is functional or constant.

## 5 Applications in genomics

In this section, we apply the fFDR method to analyze data from two studies, one in a genetics of gene expression (eQTL) study on baker’s yeast and the other in an RNA-seq differential expression analysis on two inbred mouse strains. We will provide a brief background on the studies and then present the analysis results for both data sets.

### 5.1 Background on the eQTL experiment

The experiment on baker’s yeast (*Sacchromyces cerevisiae*) has been performed by Smith and Kruglyak (2008), where genome-wide gene expression was measured in each of the 109 genotyped strains under two conditions, glucose and ethanol. Here we aim to identify genetic associations (technically, genetic linkage in this case) between pairings of expressed genes and SNPs among the samples grown on glucose. In this setting, the null hypothesis is “no association between a gene-SNP pair”, and the functional null proportion denotes the prior probability that the null hypothesis is true. The data set from this experiment is referred to as the “eQTL dataset”.

To keep the application straightforward, we consider only intra-chromosomal pairs, for which the genomic distances can be defined and are quantile normalized to give the informative variable *Z*. (Note that the fFDR framework can be used to consider all gene-SNP pairs by estimating a standard *r*(*p*) = *π*_0_/*f*(*p*) all inter-chromosomal pairs and then combining these with the estimated *r*(*p*, *z*) from the intra-chromosomal pairs to form the significance regions given in (6).) In this application, the genomic distance is calculated as the nucleotide basepair distance between the center of the gene and the SNP; the Wilcoxon test of association between gene expression and the allele at each SNP is used; and the p-values of such tests are obtained. We emphasize that the difference in the p-value histograms for the six strata shown in Figure 1a shows that a functional null proportion *π*_0_(*z*) is more appropriate.

### 5.2 Background on the RNA-seq study

A common goal in gene expression studies is to identify genes that are differentially expressed across varying biological conditions. In RNA-seq based differential expression studies, this goal is to detect genes that are differentially expressed based on counts of reads mapped to each gene. The null hypothesis is “no differential expression (for a gene) between the two conditions”, and the functional null proportion denotes the prior probability that the null hypothesis is true. For an RNA-seq study, the quantile normalized per-gene read depth is the informative variable *Z* that we utilized, which affects the power of the involved test statistics (Tarazona et al., 2011) or the prior probability of differential expression (Robinson et al., 2015).

We utilized the RNA-seq data studied in Bottomly et al. (2011), due to its availability in the ReCount database (Frazee et al., 2011) and because it had previously been examined in a comparison of differential expression methods (Soneson and Delorenzi, 2013). The dataset, referred to as the “RNA-seq dataset”, contains 102.98 million mapped RNA-seq reads in 21 individuals from two inbred mouse strains. As proposed in Law et al. (2014), we normalized the data using the `voom` R package, fitted a weighted linear least squares model to each gene expression variable, and then obtained a p-value for each gene based on a t-test of the coefficient corresponding to mouse strain.

### 5.3 Estimating the functional null proportion in the two studies

We applied to these two data sets our estimator 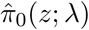 of the functional null proportion *π*_0_(*z*) utilizing the GLM, GAM, and Kernel methods. Figure 6 shows 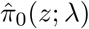 for these two data sets, and the tuning parameter *λ* has been chosen to be the one that minimizes the integrated bias and integrated variance of the function 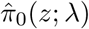; see Figure A.1 in Appendix A for details on choosing *λ*. In both data sets, 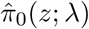 based on the GAM and Kernel methods give very similar estimates, with the one based on the GLM method more distinct, likely because the latter puts stricter constraints on the shape of *π*_0_(*z*). By comparing Figure 6 to the results in Figure 5, we see that in this RNA-seq study the read depths appear to affect the prior probability of differential expression, *π*_0_(*z*).

**Figure 5:**
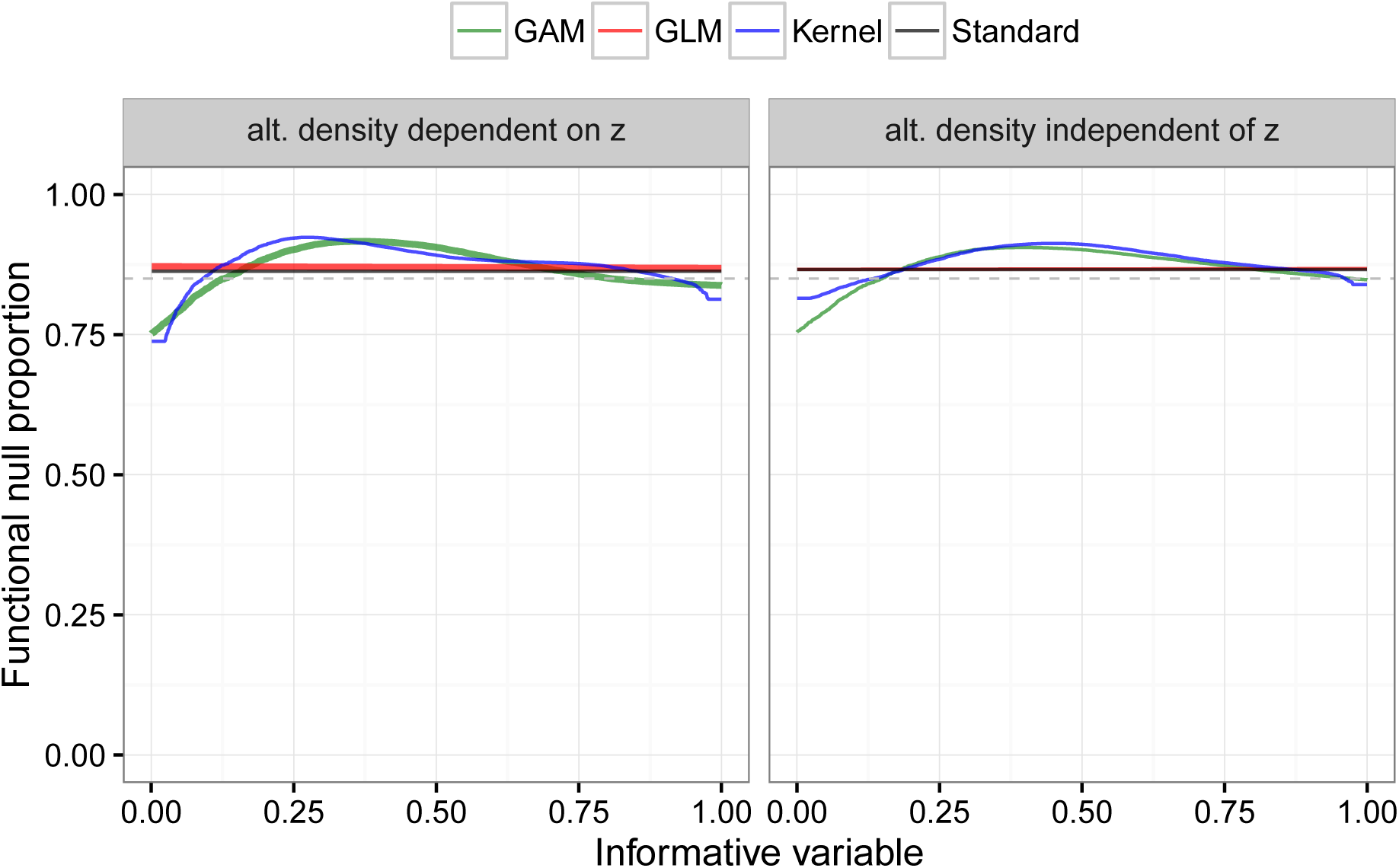
The estimator 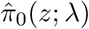 when *π*_0_ is constant. The plot titles indicate whether the alternative density *f*_1_(*p*|*z*) depends on *z* or not. In the legend “Standard” refers to the estimate of *π*_0_ provided by the `qvalue` package (Storey et al., 2015), and the dashed line indicates the true value *π*_0_ = 0.85.

As expected, the estimator 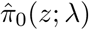 for the eQTL dataset increases with genomic distance, indicating that a distal gene-SNP association is less likely than local association. Using the GAM and Kernel methods, 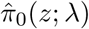 ranges from about 0.54 for very local associations and to about 0.82 for distant gene-SNP pairs. In the RNA-seq dataset, 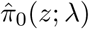 obtained by the GAM and Kernel methods decreases from around 0.97 to around 0.47 as read depth increases.

In the eQTL dataset, *λ* values from 0.4 to 0.8 led to very similar shapes of 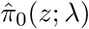 (Figure 6), and the integrated bias of 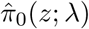 is always low compared to its integrated variance, leading to the choice of *λ* = 0.4 for all three methods. This likely indicates that the test statistics implemented in this experiment have high power, leading to low bias in estimating *π*_0_(*z*). In contrast, in the RNA-seq dataset, the integrated bias of 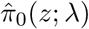 does decrease as *λ* increases, leading to a choice of a higher *λ*. While the choice of *λ* for 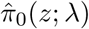 may deserve further study, it is clear from these applications that the choice of *λ* has a small effect on 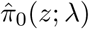 and that it is beneficial to employ a functional *π*_0_(*z*) of the informative variable rather than a constant *π*_0_.

**Figure 6:**
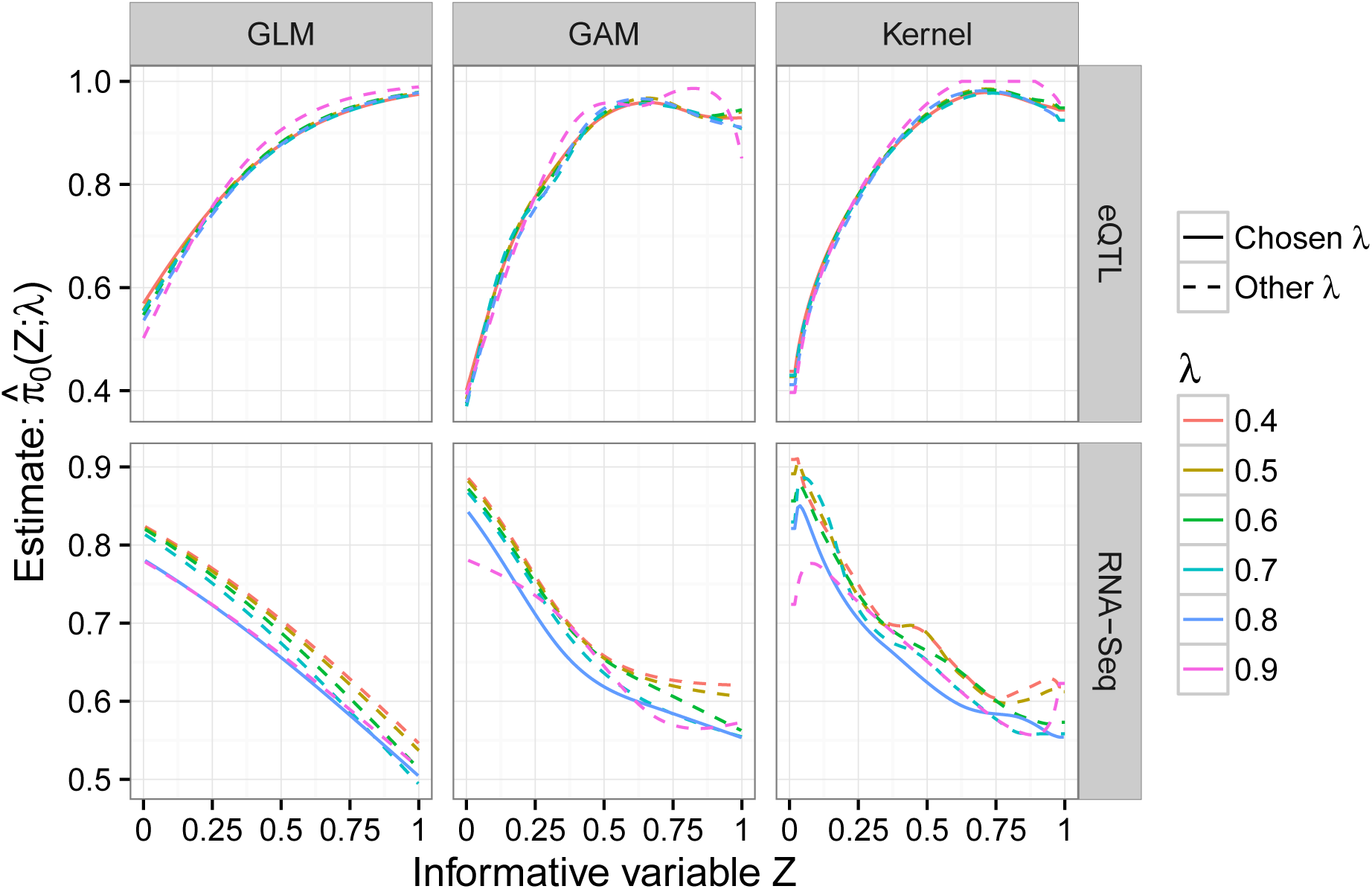
Estimate 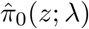 of the functional null proportion *π*_0_(*z*) for the eQTL and RNA-seq studies, using the GLM, GAM or Kernel method. Each plot shows the estimate 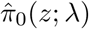 for different values of the tuning parameter *λ*, where the solid curve corresponds to the chosen tuning parameter value. The tuning parameter is chosen to balance the trade-off between the integrated bias and variance of the function 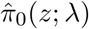; details on how to choose *λ* are given in Appendix A.

### 5.4 Application of fFDR method in the two studies

For the eQTL analysis described in Section 5.1, 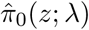 based on the GAM method is used (see Figure 6), and the fFDR method is applied to the p-values of the tests of associations and the quantile normalized genomic distances. At the target FDR of 0.05, the fFDR method found 7579 associated gene-SNP pairs, the standard FDR method 5655, and the two methods shared 5450 discoveries. Figure 7a shows that, at all target FDR levels, the fFDR method has higher power than the standard FDR method. In addition, Figure 7b reveals that the significance region of the fFDR method is greatly influenced by the gene-SNP distance, as the p-value cutoff for significance is higher for close gene-SNP pairs and lower for distant gene-SNP pairs. This, together with Figure 7c, means that some q-values *q*(*p_i_*,*z_i_*) for the fFDR method can be larger than the q-values *q*(*p_i_*) of the standard FDR method. Thus, the use of an informative variable by the fFDR method changes the significance ranking of the null hypotheses and increases the power of multiple testing at the same target FDR level.

**Figure 7:**
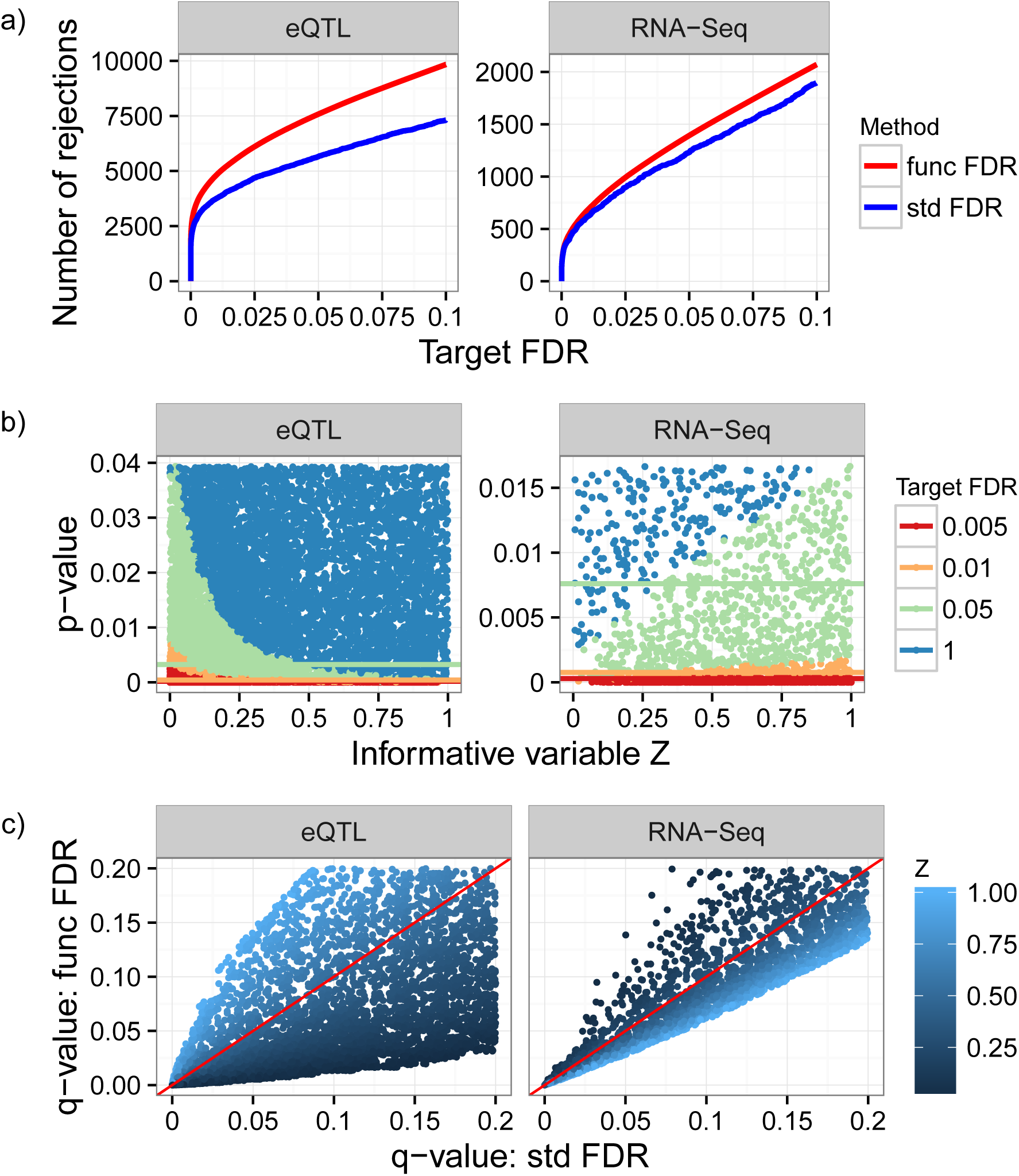
The fFDR method applied for multiple testing in the eQTL and RNA-seq analyses. a) Number of significant hypothesis tests at various target FDRs. The fFDR method (func FDR) has more significant tests than the standard FDR method (std FDR) at all target FDRs. b) The significance regions of the fFDR method for various target FDRs, indicated by scatter plots of the p-values and informative variable. The horizontal lines indicate the significance thresholds that would be used by the standard FDR method at the same target FDRs. Clearly, these lines do not take the informative variable into account. c) A scatter plot comparing the q-values for the standard FDR method (*x* axis) to the q-values for the fFDR method (*y* axis), colored based on the informative variable *Z* with reference line *x* = *y* in red. It is clear that the fFDR method re-ranks the significance of hypotheses tests.

For the RNA-seq analysis, the estimator 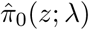 based on the GAM method is used (see Figure 6), and the fFDR method is applied to the p-values of the tests for differential expression and the quantiles of the read depths. Similar to the eQTL analysis, the fFDR method has a larger number of significant hypothesis tests at all target FDR levels; see Figure 7a. At the target FDR of 0.05, the fFDR method found 1392 genes to be differentially expressed while the standard FDR method found 1231, and the two methods shared 1202 discoveries. In this RNA-seq analysis, the fFDR method has a smaller improvement in power (see Figure 7a), the differences between the q-values for the fFDR method and those for the standard FDR method are smaller (see Figure 7c), and the significance region of the fFDR method is less affected by the informative variable *Z* (see Figure 7b). This may be because the total number of differentially expressed genes in the RNA-seq study was small or the test statistics applied in this experiment were already powerful.

## 6 Discussion

We have proposed the fFDR methodology to utilize additional information on the prior probability of a null hypothesis being true or the power of the family of test statistics in multiple testing. It employs a functional null proportion of true null hypotheses and a joint density for the p-values and informative variable. Our simulation studies have demonstrated that the fFDR methodology is more accurate and informative than the standard FDR method.

Besides the eQTL and RNA-seq analyses demonstrated here, the fFDR methodology is applicable to multiple testing in other studies. For example, it can be applied to genome-wide association studies such as those conducted in Dalmasso et al. (2008) and Roeder et al. (2006), where an informative variable can incorporate differing minor allele frequencies or information on the prior probability of a gene-SNP association obtained from previous genome linkage scans. It can also be used in brain imaging studies, e.g., those conducted or reviewed in Benjamini and Heller (2007) and Chumbley and Friston (2009), to integrate as the informative variable spatial-temporal information on the measurements for the voxels.

Finally, the fFDR methodology can be extended to the case where p-values or the status of null hypotheses are dependent on each other. In this setting, the corresponding decision rule may be different from that obtained here. On the other hand, the methodology can be extended to incorporate a vector of informative variables. This could be especially appropriate when additional information can not be compressed into a univariate informative variable.

## Software

The methods described in this paper will be made available in the `qvalue` R package, available at https://github.com/StoreyLab/qvalue (most recent version) or http://www.bioconductor.org/packages/release/bioc/html/qvalue.html (official release).

## Acknowledgement

This research was supported in part by National Institutes of Health grant R01 HG002913 and Office of Naval Research grant N00014-12-1-0764.

## Appendix

In this appendix, we show how to choose the tuning parameter *λ* for the estimator 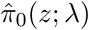 of the functional null proportion *π*_0_(*z*) and how to ensure that the estimator 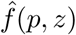 of the joint density *f*(*p*, *z*) of the p-value and informative variable (*P*, *Z*) has the same monotonicity property of *f* when evaluated at the observations 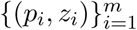

### A Choice of tuning parameter for estimators of the functional null proportion

The GLM, GAM and Kernel methods of estimating the functional null proportion *π*_0_(*z*) all rely on a tuning parameter *λ* that controls the trade-off between the bias and variance of the estimator 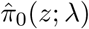. One observation we have made from the simulation studies is the following: 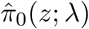 with smaller *λ* tends to capture the correct shape of *π*_0_(*z*) but with different levels of bias, while 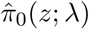 with larger *λ* may have poorly estimated the shape of *π*_0_(*z*) but with less bias, all with respect to the mean

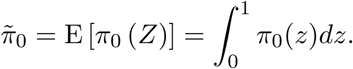

So, a simple rule of thumb is to pick a 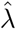 for which 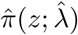 is the least biased but still has a similar shape to those of 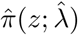 obtained with larger *λ*. In other words, the desired 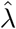 should balance the integrated bias and variance of 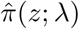. The following formalizes this intuition.

For a function *ϕ_λ_* that satisfies the constraint

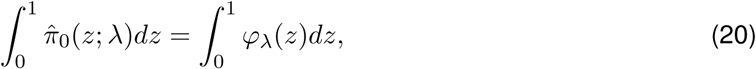
let

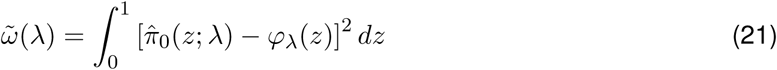
be an approximate integrated variance of 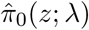. Further, let

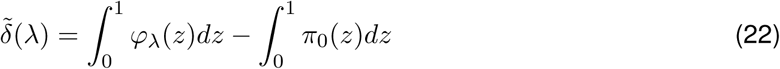
be an approximate integrated bias of 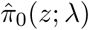. Set 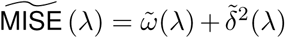. Then 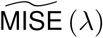 measures the “bias” and “variance” of 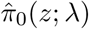 with respect to a reference estimate *ϕ_λ_*, and a 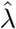 that minimizes 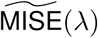 will achieve the desired trade-off without much computational cost or much loss in the accuracy in estimating *π*_0_(*z*).

With the above preparation, the procedure to determine 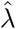 is as follows:

1. Let 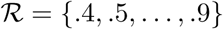 be a finite set of 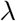 values in (0, 1). For each 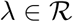, obtain 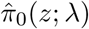 using a method provided in Section 3.1.
2. Let 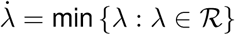. For each 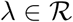, estimate the function *ϕ_λ_* as

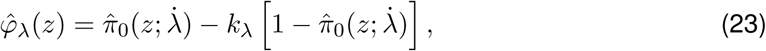
where *k_λ_* is chosen such that 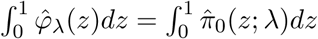. Use 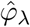 to estimate 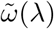 as

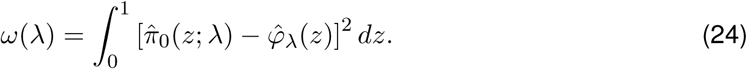
3. Use the method in Storey et al. (2004) to compute 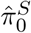 in (16) as an estimate of the mean 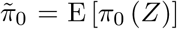; use 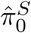 to estimate 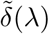 as

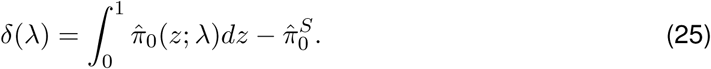
4. Set

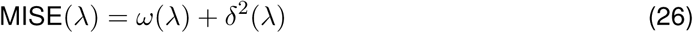
as an estimate of 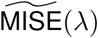, and choose 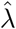 as

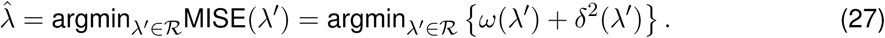 Set 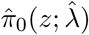 as the estimate of *π*_0_(*z*).

Figure A.1 illustrates how *λ* is chosen when estimating *π*_0_(*z*) for each of the data sets analyzed in Section 5.3.

**Figure A.1.**
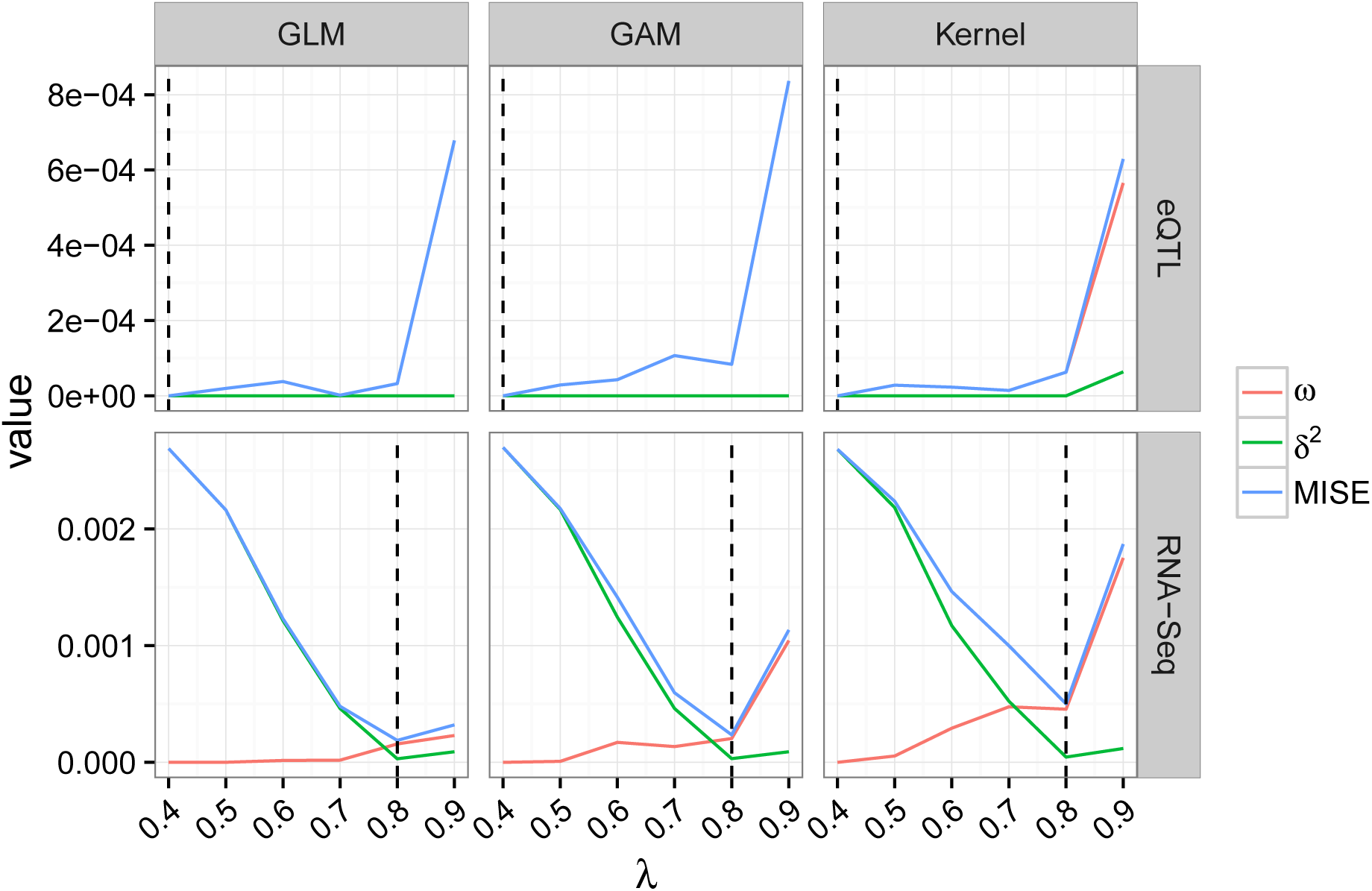
Determining the tuning parameter *λ* in the estimator 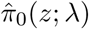 shown in Figure 6 for the eQTL and RNA-seq analyses. In the legend, *ω* is the estimated integrated variance defined by (24), *δ*^2^ the square of the estimated integrated bias defined by (25), and MISE the estimated “mean integrated squared error” defined by (26). The chosen 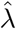 minimizes the MISE and is indicated by the vertical dashed line.

### B Ensuring a monotonicity property of the estimator of the joint density

Recall that the joint density for the p-value and informative variable (*P*, *Z*) satisfies

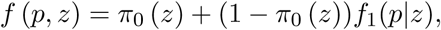
where *f*_1_(*p*|*z*) is the conditional density of the p-value under the true alternative hypothesis given *Z* = *z*. If *f*_1_(*p*|*z*) is non-increasing in *p* for each fixed *z*, then so is *f*(*p*, *z*) and so should be its estimate 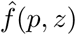. In order to make 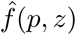 have such a monotonicity property when evaluated at the observations {(*p_i_*,*z_i_*), 1 ≤ *i* ≤ *m*}, we modify the estimate 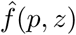 of *f*(*p*, *z*) discussed in Section 3.2 as follows. For each *i* = 1,…, *m*, set

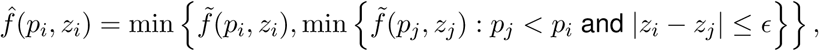
where *ϵ* (by default being 0.02 in this paper) is chosen to be small and positive because *Z* ~ Uniform(0, 1) and Pr(*z_i_* = *z_j_*)=0 for *i* ≠ *j*. This guarantees that, for all pairs (*p_i_,z_i_*), (*p_j_,z_j_*) ∈ [0, 1]^2^,

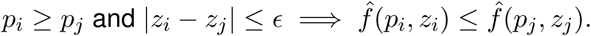

In the unusual event that *f*(*p*, *z*) is non-decreasing in *p* for each fixed *z*, then an analogous algorithm can be performed satisfying the non-decreasing property.

## Footnotes

1 SNP stands for “single nucleotide polymorphism” and is a common type of genetic marker utilized in eQTL studies.

